# Evolutionary landscape of mosquito viral integrations

**DOI:** 10.1101/385666

**Authors:** Elisa Pischedda, Francesca Scolari, Federica Valerio, Rebeca Carballar-Lejarazú, Paolo Luigi Catapano, Robert M. Waterhouse, Mariangela Bonizzoni

**Author notes:** Corresponding author: Mariangela Bonizzoni. Research was conducted at the Department of Biology and Biotechnology of the University of Pavia, Italy.

## Abstract

The sequenced genome of the arboviral vector mosquito, *Aedes albopictus,* is replete with repetitive DNA and it harbors an unusually large number of endogenous viral sequences, collectively called Nonretroviral Integrated RNA Virus Sequences (NIRVS). NIRVS are enriched both within protein-coding gene exons and PIWI-interacting RNA (piRNA) clusters, where they encode piRNAs. Based on these features, NIRVS have been proposed to function as novel mosquito antiviral immune factors. However, the relative importance and contributions of different NIRVS as functional antiviral elements and their mechanisms of action remain open questions.

We apply an analytical approach that intersects computational, evolutionary and molecular methods to identify NIRVS most likely affecting mosquito immunity. Using this strategy, we show that NIRVS are a highly dynamic component of the *Ae. albopictus* repeatome, which nevertheless maintains a core set of seemingly the oldest NIRVS with similarity to Rhabdoviruses. Population-level polymorphism of NIRVS varies depending on whether they occur in intergenic regions, piRNA clusters or are part of gene exons. NIRVS from piRNA clusters are differentially widespread in diverse populations but conserved at the sequence level. This is consistent with the hypothesis that they act analogously to fragments of transposable elements in piRNA clusters and contribute to piRNA-based immunity. Among NIRVS from gene exons, AlbRha52 and AlbRha12 have the hallmarks of domestication as they are fixed across populations, stably expressed, and as polymorphic at the sequence level as fast-evolving genes. Overall these results support the hypothesis that NIRVS contribute to mosquito immunity, potentially through diverse modes of action.

## Introduction

The amount and the type of repeated DNA sequences, collectively called the “repeatome”, affect the size, organization and evolution of eukaryotic genomes (Maumus and Quesneville 2014). Transposable elements (TEs) are the major and most-studied components of the repeatome because of their potential mutagenic effects (Gilbert and Feschotte 2018). TEs evolve through a “burst and decay model” whereby newly acquired TEs can multiply rapidly in a genome. The “burst” phase is followed by low amplification periods, the “decay” moments, when TEs tend to accumulate mutations and become inactive (Maumus and Quesneville 2016). In eukaryotes, TE mobilization during germline formation is counterbalanced by the activity of the PIWI-interacting RNA (piRNA) pathway, the most recently identified of three small RNA-based silencing mechanisms (Brennecke et al. 2007; Guzzardo et al. 2013; Gainetdinov et al. 2017). Briefly, Argonaute proteins of the PIWI-subfamily associate with small RNAs of 25-30 nucleotides, called PIWI-interacting RNAs (piRNAs), and together they silence TEs based on sequence complementarity (Toth et al. 2016). piRNAs arise from genomic regions called piRNA clusters, which contain fragmented sequences of previously acquired TEs. piRNA clusters are dynamic components of the *Drosophila melanogaster* genome and their structural composition has been linked to differential regulatory abilities. For instance, the ability of the *D. melanogaster* master piRNA locus *flamenco* to control transposons such as *gypsy*, *ZAM* and */defix* was shown to be dependent on frequent chromosomal rearrangements, loss or gain of fragmented TE sequences (Zanni et al. 2013; Guida et al. 2016). Additionally, variations in the composition of subtelomeric piRNA clusters were observed upon adaptation to laboratory conditions of *D. melanogaster* wild collected flies (Asif-Laidin et al. 2017). Importantly, structural differences in subtelomeric piRNA clusters did not impair host genome integrity and occurred with the maintenance of conserved groups of sequences, which could be alternatively distributed among different strains (Asif-Laidin et al. 2017).

Unexpectedly, besides TE fragments, piRNA clusters contain sequences from nonretroviral RNA viruses, which produce piRNAs, in the genome of arboviral vectors like the mosquitoes *Aedes aegypti* and *Aedes albopictus* (Arensburger et al 2011; Olson and Bonizzoni 2017; Palatini et al. 2017; Whitfield et al. 2017). This observation is in line with recent experimental evidence that extend the role of the piRNA pathway to immunity against viruses in *Aedes* mosquitoes, differently than in *D. melanogaster* (Petit et al. 2016; Miesen et al. 2016) and show that piRNAs from integrated viral sequences are differentially expressed following viral challenge of *Ae. albopictus* (Wang et al. 2018). As such Nonretroviral Integrated RNA Virus Sequences (NIRVS) have been proposed as novel immunity factors of arboviral vectors (Palatini et al. 2017; Olson and Bonizzoni 2017; Whitfield et al. 2017). However, the organization, stability and mode of action of NIRVS in mosquito genomes are poorly understood.

The landscape of viral integrations in the genome of *Ae. aegypti* and *Ae. albopictus* mosquitoes is quite complex. *Aedes* species are rare examples within the animal kingdom because they harbor dozens of NIRVS from different viruses, such as Flaviviruses and Mononegavirales, primarily Rhabdoviruses and poorly characterized Chuviruses (Katzourakis and Gifford 2010; Fort et al. 2012; Palatini et al. 2017; Whitfield et al. 2017). In all other animals in which NIRVS have been identified, including mammals, birds and ticks, NIRVS appear to be mainly virus-host specific and tend to be found in low numbers (<20) (Katzourakis and Gifford 2010; Belyi et al. 2010; Holmes 2011; Kryukov et al. 2018). NIRVS identified in the *Ae. aegypti* and *Ae. albopictus* genomes are not homologous, indicating independent integration events. However, NIRVS of both species encompass fragmented viral open reading frames (ORFs) and are enriched within piRNA clusters. In *Ae. albopictus* NIRVS are also enriched within coding sequences (Palatini et al. 2017). Taken together these findings support the hypothesis that NIRVS contribute to host biology. However, whether all NIRVS or some are functionally active elements and how they work are still open questions.

We postulate that given the abundance and the sequence similarity of NIRVS of the *Ae. albopictus* genome, an ideal strategy to identify those viral integrations that most likely are indispensable functional genomic elements is to investigate the genomic architecture of NIRVS in an evolutionary framework. If NIRVS act similarly to TE fragments within piRNA clusters, their overall genomic landscape should be variable across host genomes, but, when NIRVS are maintained within a genome, their sequence should be conserved (Zanni et al. 2013; Goriaux et al. 2014; Asif-Laidin et al. 2017). If NIRVS within coding genes have been co-opted for host functions, they are expected to be expressed and if they are involved in immunity functions, they are expected to be under rapid evolution because their products should act against fast evolving viruses (Frank and Feschotte 2017). Finally, if NIRVS result from fortuitous events and are viral fossils, they should have reached fixation and evolve at a neutral rate (Aswad and Katzourakis 2012; Katzourakis 2013).

Under these premises, we studied the distribution of NIRVS in the genome of *Ae. albopictus* geographic samples and the sequence polymorphism of NIRVS in relation to fast-and slow-evolving mosquito genes. Using an analytical approach that intersects computational, evolutionary, and molecular approaches we show that NIRVS are a dynamic component of the *Ae. albopictus* repeatome. The landscape of NIRVS is variable among geographic samples and their levels of sequence polymorphism is heterogeneous. NIRVS with similarities to Rhabdoviruses appear more widespread and older integrations than those with similarities to Flaviviruses. NIRVS annotated in intergenic regions appear more variable than those mapping within piRNA clusters or gene exons. Among NIRVS identified within gene exons, six are fixed and stably expressed, albeit showing different levels of polymorphism. Overall these results support the hypothesis that NIRVS may contribute to mosquito immunity and highlight their diverse modes of action.

## Results and Discussion

### Whole Genome Sequencing (WGS) data and molecular approaches are concordant in showing that viral integrations are abundant in the genome of *Ae. albopictus*

We first validated the use of a WGS-based approach to study the landscape and the polymorphism of viral integrations within the *Ae. albopictus* genome given the abundance (*i.e.* 72 loci) and the sequence-similarity of NIRVS that were bioinformatically characterized in the Foshan reference strain (Palatini et al. 2017). For this task, we focused on NIRVS with similarities to Flaviviruses and we compared estimates of the copy number (CN) of each NIRVS based on bioinformatic and molecular approaches (*i.e.* qPCR and Southern Blotting experiments) (table 1). WGS data were obtained from 16 mosquitoes (eight males and eight females) whose genomes were individually-sequenced (*i.e.* singly-sequenced mosquitoes or SSMs) after having forced them in single mating and collected their progeny. qPCR data were obtained from the SSM progeny and Southern Blotting data were derived from pools of Foshan mosquitoes. As shown in table 1, CN estimates of NIRVS derived from WGS data and molecular experiments are consistent, with few exceptions. For instance, despite the appearance of one band in hybridization experiments (supplementary fig. S1), we hypothesized that qPCR-based CN values below 1 for AlbFlavi4 were due to its presence in hemizygosity in some mosquitoes. We verified that hemizygosity of NIRVS gives rise to qPCR-based CN values below 1 by quantifying AlbFlavi4 in the progeny of a family in which the mother and the father bioinformatically showed presence and absence of this locus, respectively (fig. 1A). The discrepancies in CN estimates for the sequence-similarity group including AlbFlavi18, AlbFlavi20 and AlbFlavi28 are explained by the different distribution of these integrations in SSMs and their occurrence in hemizygosity in some individuals (fig. 1B). AlbFlavi20 and AlbFlavi18 are rare: they were found at a frequency of 0.13 and 0.25, respectively. This is in contrast to AlbFlavi28, which was detected at a frequency of 0.69 in SSMs and, when present in hemizygosity, gives CN values below one in qPCR experiments (fig. 1B). Higher CN values from Southern blot analyses with respect to bioinformatic data are due to the polymorphism of these F-NIRVS in mosquitoes (fig. 1C) and their high sequence similarity that limited the design of NIRVS-specific probes (Supplementary Material online). Overall, these results support the application of WGS, coupled with bioinformatic analyses, to study the genomic architecture of NIRVS.

**Table 1:** Comparison among WGS data, qPCR>based copy number (CN) and Southern Blot (SB) results of F>NIRVS. qPCR-based CN was estimated for five Foshan mosquitoes. SB data were generated hybridizing F-NIRVS specific probes to pooled DNA from 10-20 mosquitoes. qPCR amplification results between 0.25-0.75 are assumed to correspond to heterozygous individuals.

| Target F-NIRVS | Exp. CN<sup>a</sup> | WGS-based frequency<sup>b</sup> | qPCR-based CN<sup>c</sup> | SB-based CN |
| --- | --- | --- | --- | --- |
| AlbFlavi19, 33 | 2 | AlbFlavi19:0; AlbFlavi33:0 | 0 | 0 |
| AlbFlavi24, 33 | 2 | AlbFlavi24:0.75; AlbFlavi33:0 | 0.57<sup>d</sup> | 1 |
| AlbFlavi31, 32, 34 | 3 | AlbFlavi31:0; AlbFlavi32:0; AlbFlavi34:1 | 0.48 | 1 |
| AlbFlavi18, 20, 28 | 3 | AlbFlavi18:0.25; AlbFlavi20:0.13 | 0.52 | 2 |
|  |  | AlbFlavi28:0.69 |  |  |
| AlbFlavi26, 37, 42 | 3 | AlbFlavi26:0.94; AlbFlavi37:0.88 | 0.88<sup>e</sup> | 3<sup>f</sup> |
|  |  | AlbFlavi42:0.69 |  |  |
| AlbFlavi22, 23, 27, 26, 37 | 5 | AlbFlavi22:0.94; AlbFlavi23:0.81; AlbFlavi27:0.88; | 1.48 | 3<sup>f</sup> |
|  |  | AlbFlavi26:0.94; AlbFlavi37:0.88 |  |  |
| AlbFlavi22, 25, 27, 26, 37 | 5 | AlbFlavi25:1 | 3.83 | 3 |
| AlbFlavi8-41 | 2 | AlbFlavi8-41:0.25 | 1.18<sup>d</sup> | 4<sup>g</sup> |
| AlbFlavi3 | 1 | AlbFlavi3:0.94 | 0.40 | 2<sup>g</sup> |
| AlbFlavi4 | 1 | AlbFlavi4:0.88 | 0.56 | 1 |
| AlbFlavi6 | 1 | AlbFlavi6:1 | 1.03 | smear<sup>h</sup> |
| AlbFlavi7 | 1 | AlbFlavi7:1 | 0.76 | 0<sup>i</sup> |
| AlbFlavi10 | 1 | AlbFlavi10:0.81 | 0.39 | 2 |
| AlbFlavi36 | 1 | AlbFlavi36:0.81 | 0.58 | 2<sup>g</sup> |
| AlbFlavi2 | 1 | AlbFlavi2:0.94 | 0.90 | 2 |
| AlbFlavi1, AlbFlavi12-17 | 2 | AlbFlavi1:0.94 | 2.40 | 2<sup>j</sup> |
| AlbFlavi12-17 | 1 | AlbFlavi12-17:0.88 | 0.69 |  |

<sup>a</sup>expected CN based on the current AaloF1 genome assembly; <sup>b</sup>frequency was calculated based on number of individuals out of the 16 singly-sequenced Foshan mosquitoes with read coverage supporting the indicated F-NIRVS (fig. 2B); <sup>c</sup>mean CN was calculated excluding individuals lacking the corresponding viral integration; <sup>d</sup>out of the tested 5 individuals, 4 gave no amplification; <sup>e</sup>qPCR primers amplified only AlbFlavi37 and AlbFlavi42. We assumed qPCR data indicate the presence of only one of the three NIRVS or two, each in a heterozygous state; <sup>f</sup>SB data were obtained with the same probe (Supplementary Material online); <sup>g</sup>see Supplementary Material online for probe hybridization features; <sup>h</sup>great variability was observed for this locus; <sup>i</sup>the high variability of the flanking regions did not allow the design of a probe of appropriate length; <sup>j</sup>SB data were obtained with the same probe (Supplementary Material online).

**Fig. 1:**
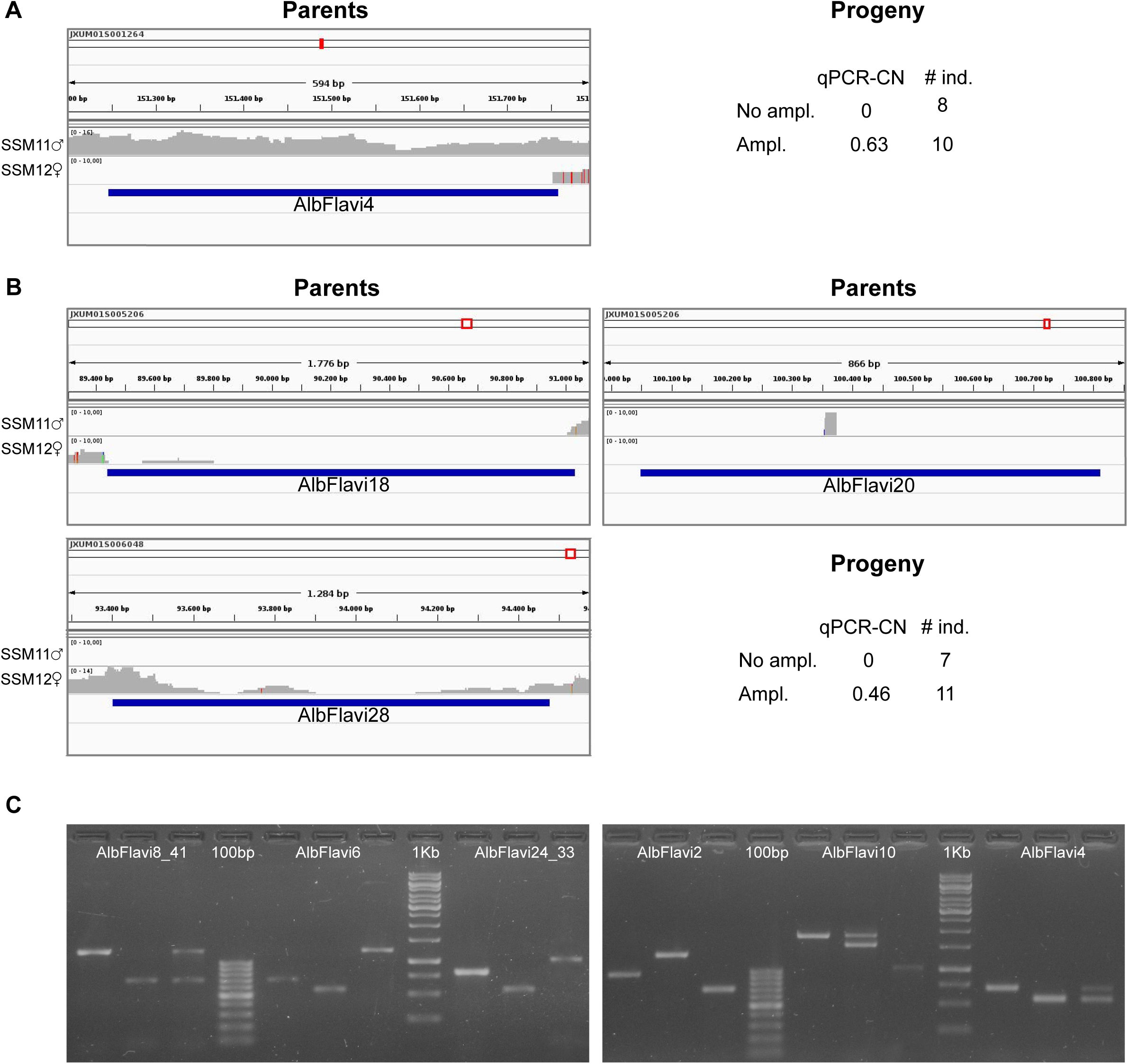
Bioinformatic and molecular approaches are concordant in identifying NIRVS. IGV visualization of the read-coverage of SSM11 and SSM12 in *(A)* AlbFlavi4 and *(B)* AlbFlavi18, AlbFlavi20 and AlbFlavi28. The progeny resulting from the mating between SSM11 and SSM12 was collected and 18 individuals were tested by qPCR to estimate F-NIRVS copy number. In AlbFlavi4, no amplification was observed for eight individuals, while ten gave results below 1, indicating hemizygosity, as expected based on the genotype of the parents. A similar situation was observed for the similarity group including AlbFlavi18, AlbFlavi20 and AlbFlavi28. *(C)* Amplification of AlbFlavi8-41, AlbFlavi6, AlbFlavi24-33, AlbFlavi2, AlbFlavi10 and AlbFlavi4 from DNA of three mosquitoes showed bands of different lengths, indicating variability of NIRVS in individual mosquitoes.

### NIRVS are variably distributed in SSMs

Given the confidence in using WGS data to identify and characterize NIRVS, we designed a pipeline to evaluate the genomic landscape of NIRVS and their polymorphism. Overall, mean base pairs (bp) occupied by NIRVS with similarities to Flaviviruses (F-NIRVS) and Rhabdoviruses (R-NIRVS) are 12095 and 19293 bp, respectively (fig. 2A). No read coverage was observed in any of the 16 SSMs for eleven NIRVS (*i.e.* AlbFlavi19, AlbFlavi31, AlbFlavi32, AlbFlavi33, AlbFlavi38, AlbFlavi39, AlbFlavi40, AlbRha43, AlbRha79, AlbRha80, AlbRha95). A total of 20 NIRVS were found in all SMMs, with a statistical enrichment for R-NIRVS (Hypergeometric test, p= 0.022) and NIRVS mapping in gene exons (Hypergeometric test, p=0.006) (fig. 2B). This “core” of 20 NIRVS included all R-NIRVS identified within the coding sequence of genes (*i.e.* AlbRha12, AlbRha15, AlbRha18, AlbRha28, AlbRha52, AlbRha85 and AlbRha9) and piRNA clusters (*i.e.* AlbRha14 and AlbRha36). Conversely, F-NIRVS were variably distributed among SSMs. Of note is AlbFlavi4, a 512bp sequence with similarity to the capsid gene of *Aedes flavivirus* (Palatini et al. 2017). AlbFlavi4 is annotated within the second exon of AALF003313 and is also included in piRNA cluster 95 (Liu et al. 2016). AlbFlavi4 produces vepi4730383, a piRNA that is upregulated upon dengue infection (Wang et al. 2018). In SSMs and *Ae. albopictus* geographic samples, variants were identified for AALF003313, only one of which includes AlbFlavi4 (fig. 2C-D).

**Fig. 2:**
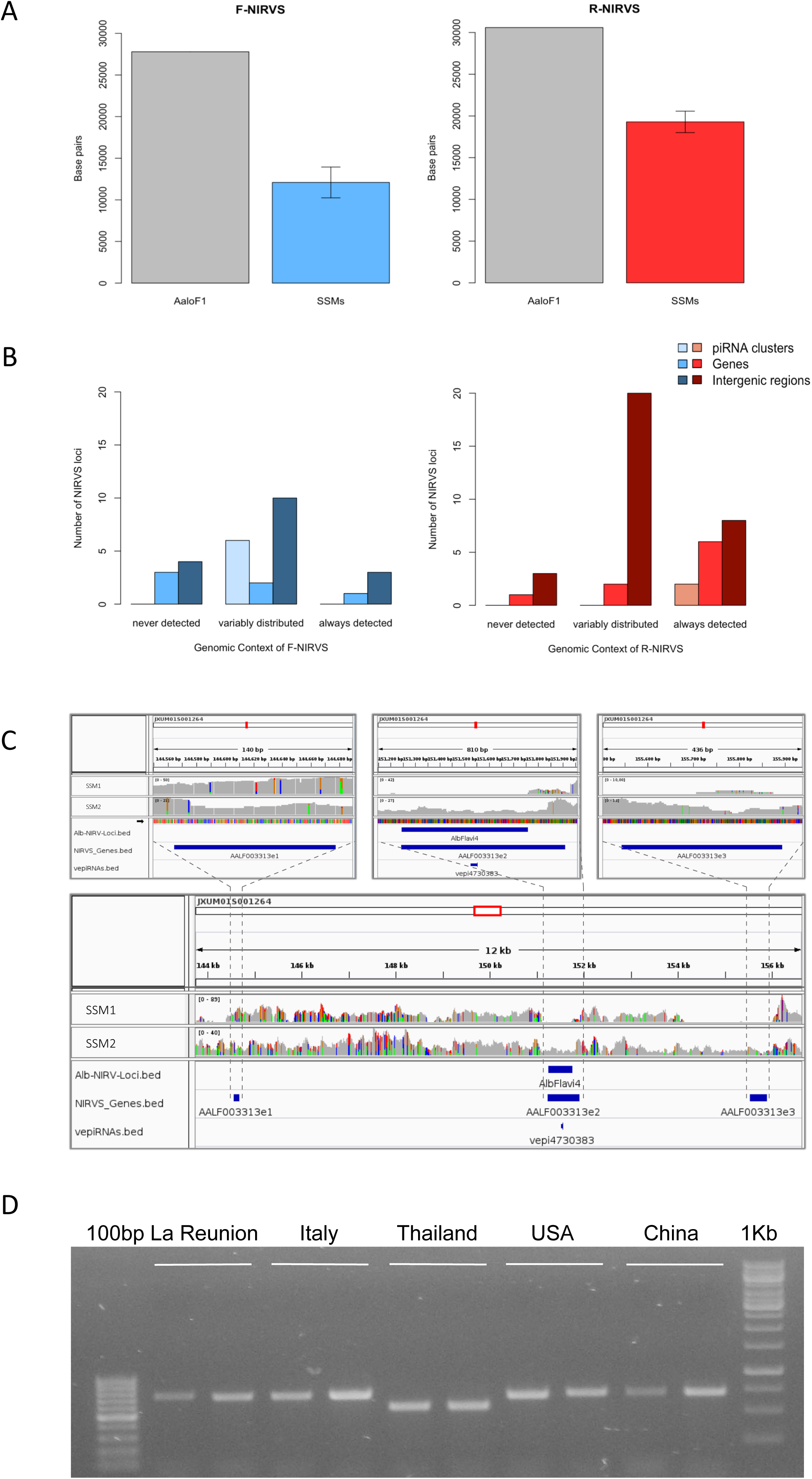
NIRVS are variably distributed in SSMs. *(A)* Flavivirus (F-NIRVS) and Rhabdovirus (R-NIRVS) loci occupancy in the 16 singly-sequenced mosquitoes (SSMs) of the Foshan strain is about half of that expected based on the annotated sequences of the reference genome assembly (AaloF1). *(B)* Number of NIRVS mapping within genes, piRNA clusters or intergenic regions, classified on the basis of read-coverage across SSMs. In A and B panels, F-NIRVS are in blue, R-NIRVS are in red. *(C)* IGV screen shot showing read-coverage at AALF003313 in SSM1 and SSM2. Positions of the three AALF003313 exons, AlbFlavi4 and vepi4730383 are indicated by blue bars. *(D)* PCR amplification of AALF003313 exon2 in ten *Ae. albopictus* geographic samples.

Together, these results demonstrate that, with an average genome occupancy of 31389 bp, NIRVS represent quantitatively a limited fraction of the mosquito repeatome. However, the enrichment of NIRVS in piRNA clusters (Palatini et al. 2017) and the fact that the pattern of NIRVS is variable in host genomes make them an important component of the repeatome. Besides being consistent with the hypothesis that NIRVS behave analogously to TE fragments within piRNA clusters, the variable distribution of NIRVS may be exploited for phylogenetic studies. Interestingly, R-NIRVS appear more widespread than F-NIRVS. This difference is intriguing considering that Mononegavirales are evolutionary more recent than *Flaviviridae* (Koonin et al. 2015). The promiscuous nature of Rhabdoviruses, which switch among hosts more frequently than any other RNA virus (Geoghegan et al. 2017), favor their wide spread, could favor their integration within host genomes, and could explain these observations.

### NIRVS as molecular markers

To test if NIRVS can be used as molecular markers in phylogeographic studies, we choose seven F-NIRVS (AlbFlavi2, AlbFlavi4, AlbFlavi8-41, AlbFlavi10, AlbFlavi36, AlbFlavi1 and AlbFlavi12-17) and six R-NIRVS (AlbRha1, AlbRha7, AlbRha14, AlbRha36, AlbRha52, AlbRha85) based on their unique occurrence in different regions of the mosquito genome and their similarity to various viral ORFs. We tested both their presence and their sequence variability in native, old and new *Ae. albopictus* populations (Manni et al. 2017). Alleles of NIRVS were differentially distributed across geographic populations so that a tree built from a matrix of shared-allele distances (DAS) proved able to differentiate mosquito populations in accordance with the historical records of *Ae. albopictus* invasive process when considering all thirteen NIRVS, only F-NIRVS or NIRVS mapping in intergenic regions (fig. 3A-C). On the contrary, when data from exclusively R-NIRVS or NIRVS identified in piRNA clusters, were analyzed, bootstrap values differentiating populations were below 50% (fig. 3D-E). This result agrees with the observation that R-NIRVS are much more widespread and persistent than F-NIRVS.

**Fig. 3:**
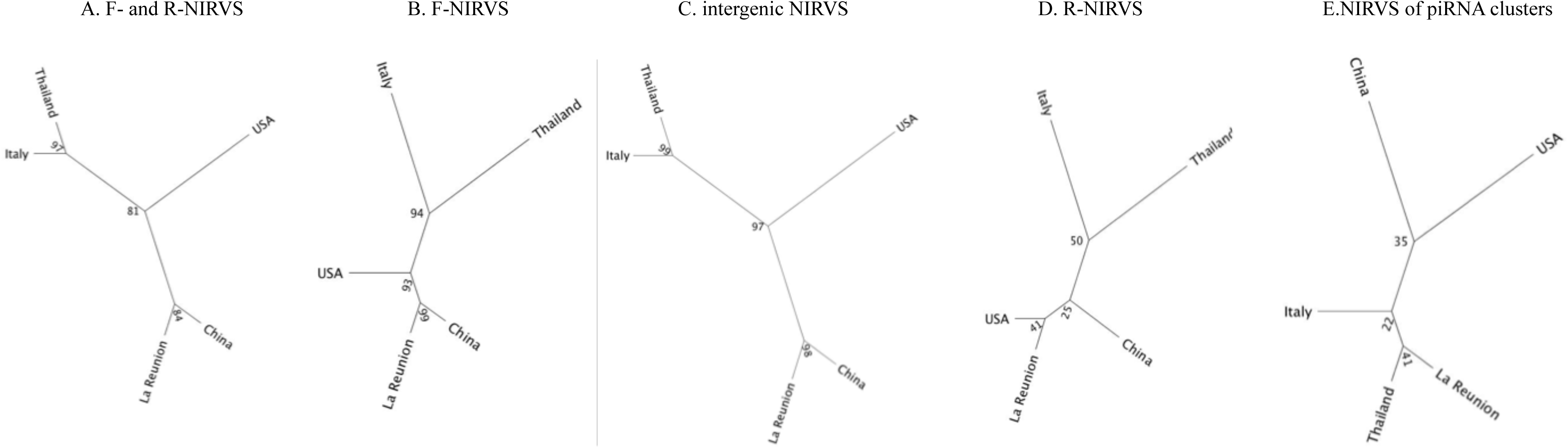
Phylogeographic distribution of NIRVS. Genetic relationships among five mosquito populations shown by a Neighbor-joining trees based on shared-allele distance using data from all 13 NIRVS that were genotyped *(A)*, only F-NIRVS *(B)* NIRVS mapping in intergenic regions *(C)*, R-NIRVS *(D)* and NIRVS mapping in piRNA clusters *(E)*. Bootstrap values are shown at the corresponding nodes.

The variable landscape of NIRVS across geographic populations should be interpreted with caution.

*Aedes albopictus* is known to be an aggressive invasive species that quickly moved out of its home range in South East Asia and reached a global distribution in the past 50 years (Bonizzoni et al. 2013; Kraemer et al. 2015). This rapid process was human-mediated and occurred through the movement of propagules (Manni et al. 2017), creating a situation of genetic admixture. Mosquito populations from newly invaded areas, such as Italy and USA, lack isolation by distance and appear genetically mixed (Manni et al. 2017; Maynard et al. 2017; Kotsakiozi et al. 2017). The occurrence of frequent bottlenecks followed by interbreeding can partly explain the variable NIRVS landscape observed here. However, the statistically significant enrichment for R-NIRVS and NIRVS mapping in exons among the “core” NIRVS indicates that selection may have also contributed in the differential distribution of NIRVS, thus emphasizing the importance of studying NIRVS in relation to their genomic context. Taken together, these results support the application of NIRVS as molecular markers for rapid species identification, based on PCR of the most widespread R-NIRVS, in cases of interceptions of immature stages or morphologically-damaged specimens. However, for the assignment of mosquitoes to a geographic region, microsatellites or ddRAD sequence markers should be used because they harbor more phylogeographic information than NIRVS (Manni et al. 2017; Kotsakiozi et al. 2017). The rapid interception of *Ae. albopictus* and the identification of its invasion routes are important tasks because of the public health relevance of *Ae. albopictus*, which is a competent vector for arboviruses such as chikungunya (CHIK), dengue (DEN), yellow fever and Zika viruses (Bonizzoni et al. 2013).

### R-NIRVS are more widespread and appear to be older integrations than F-NIRVS

The wider distribution of R-NIRVS with respect to F-NIRVS suggests R-NIRVS are older integrations. To verify this hypothesis, we tested the genealogy of R-NIRVS and F-NIRVS in comparison to circulating Rhabdoviruses and Flaviviruses. Relative timetrees were generated for i) F-NIRVS and corresponding NS3 and NS5 viral proteins from representative Flaviviruses, and ii) R-NIRVS and corresponding L and G proteins of representative Rhabdoviruses. Timetrees showed shorter divergence times between F-NIRVS and viral proteins than R-NIRVS and viral proteins. This clearly indicated multiple integration events (fig. 4A-D). Estimates of integration times were calculated for those NIRVS, whose variability was analyzed across five geographic populations, assuming comparable mutation rates between *Ae. albopictus* and *D. melanogaster* genes, that is 3.5-8.4×10^-9^ per site per generation (Haag-Liautard et al. 2007; Keightley et al. 2009), and a range of generations per year between 6 and 16, accounting for mosquitoes from temperate and tropical environments, respectively (Manni et al. 2017). Under these conditions, R-NIRVS integrated between 36 thousand and 2.9 million years ago (mya) and F-NIRVS between 6.5 thousand and 2.5 mya (fig. 5). This large window supports the conclusion that integration of viral sequence is a dynamic process occurring occasionally at different times. As shown in fig. 5, estimates of integration times varied greatly depending on the genomic context of NIRVS. NIRVS annotated within gene exons appear statistically more recent than NIRVS of piRNA clusters (ANOVA, p<0.001). Besides reflecting a different integration time, this result is consistent with the hypothesis that integrations within exons are under rapid evolution, a hallmark of domestication (Frank and Feschotte 2017). Selection constraints on sequences within piRNA clusters have been previously identified in both flies and mice (Chirn et al. 2015). This is despite piRNAs having an incredible sequence diversity and their biogenesis and processing not being linked to any common sequence or structural motifs (Huang et al. 2017).

**Fig. 4:**
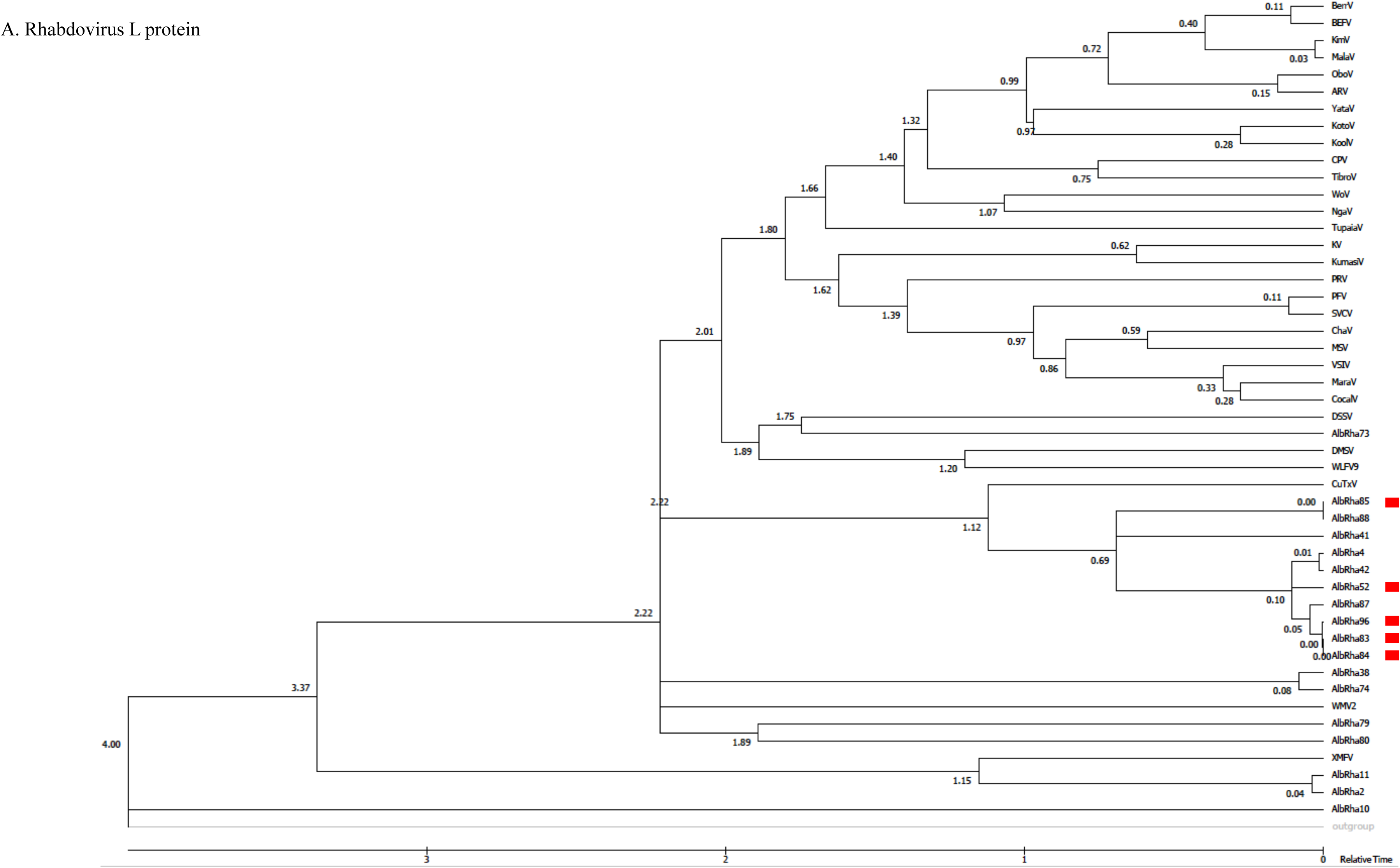
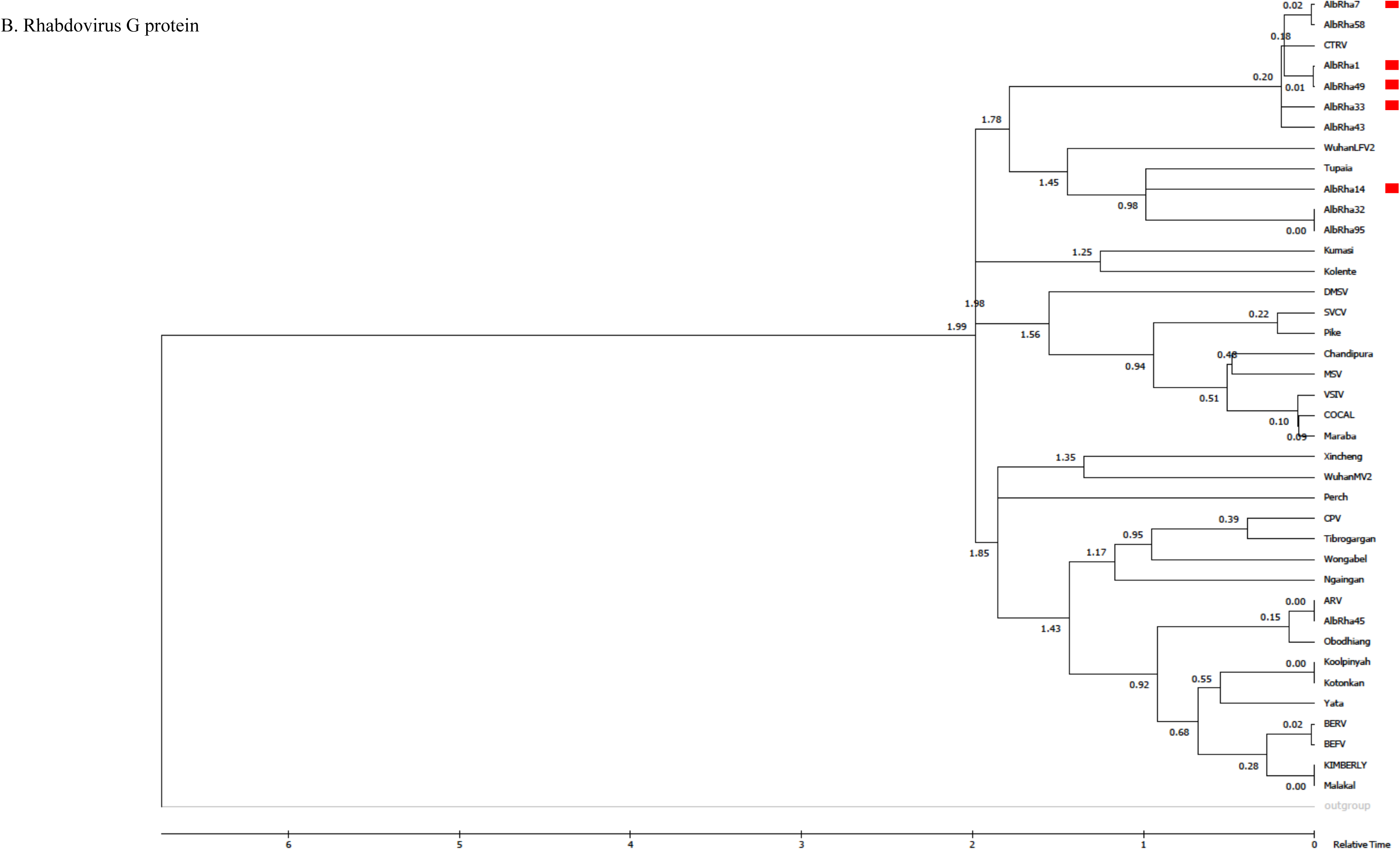
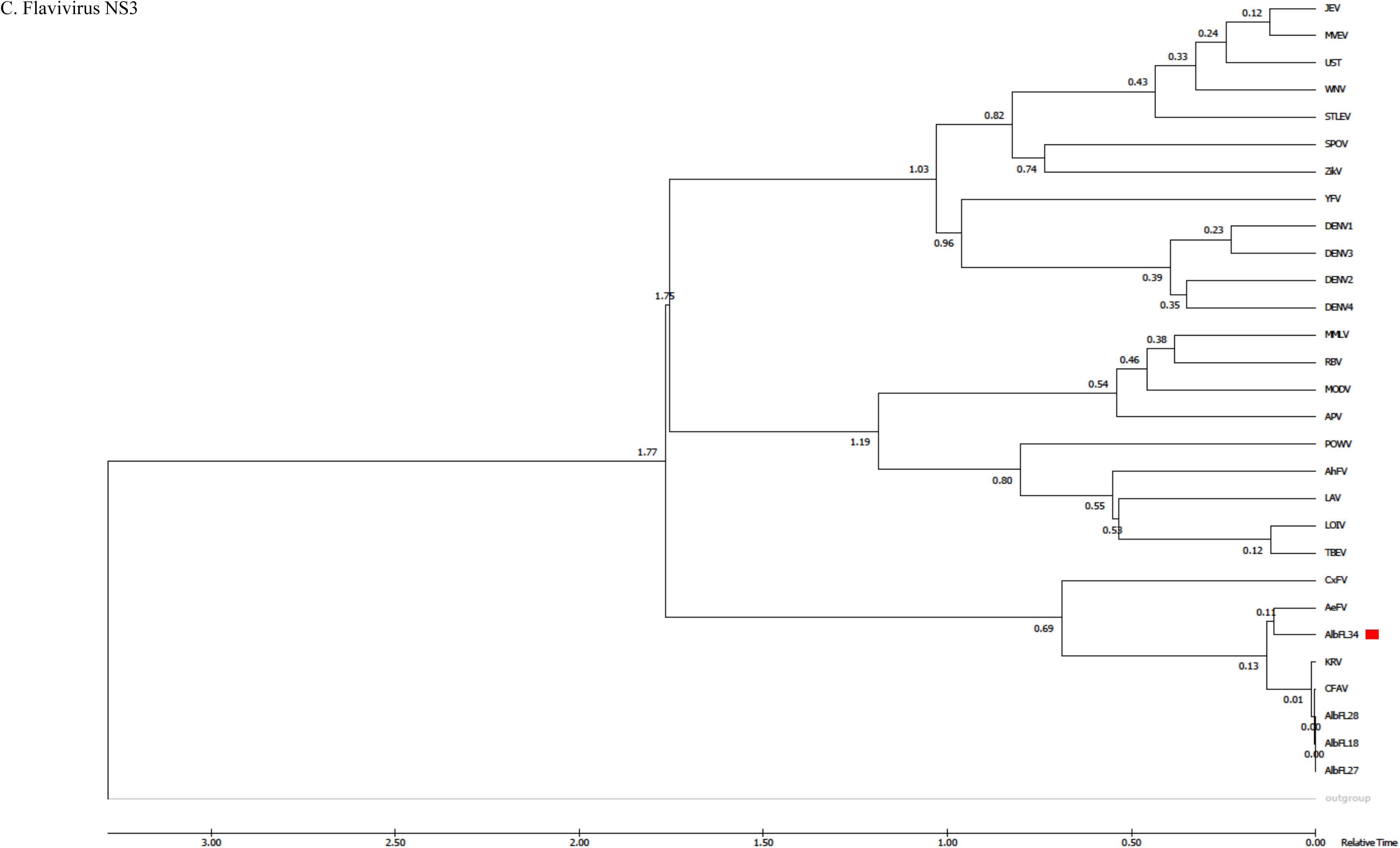
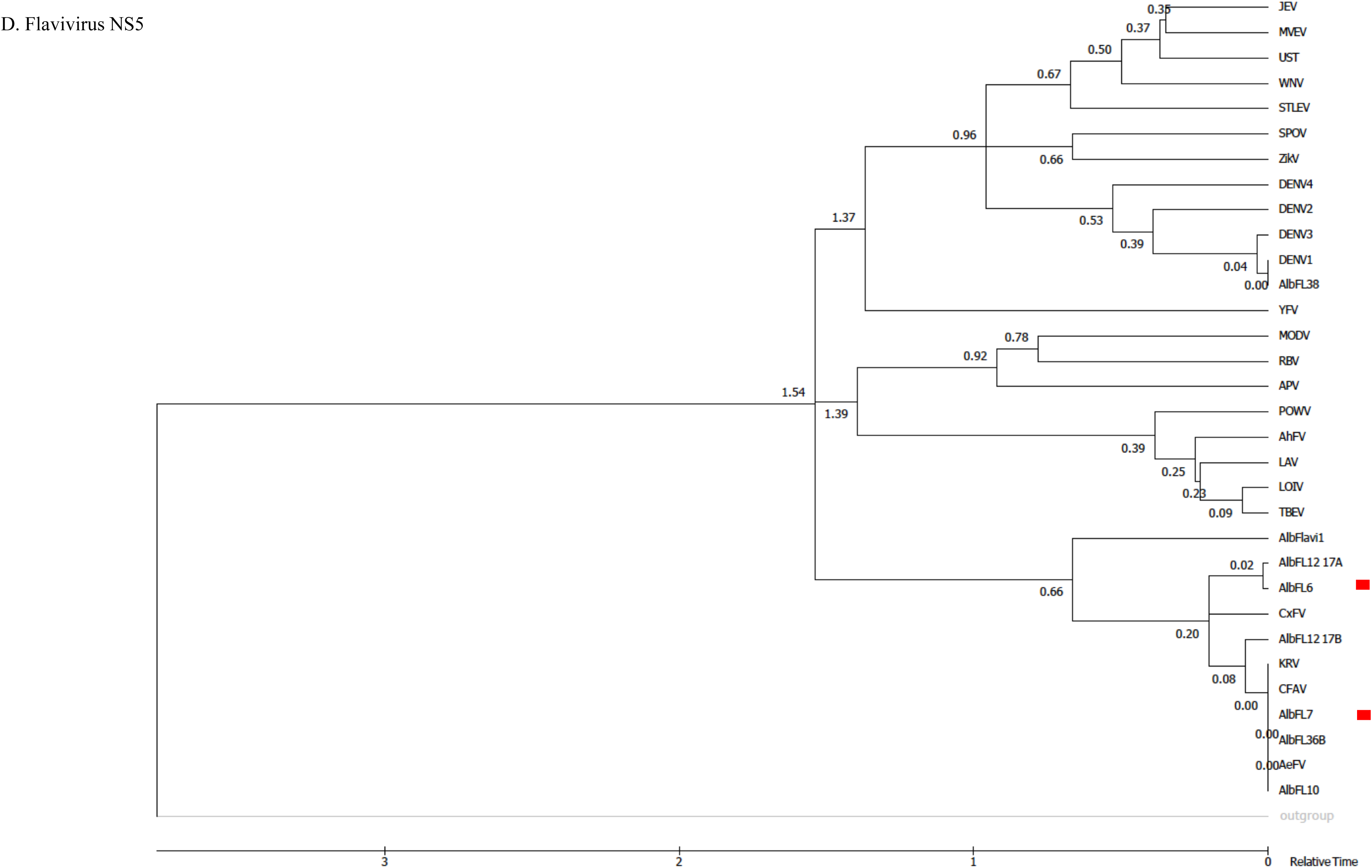
Phylogenetic analyses of FN- and R-NIRVS. *(A)* Molecular phylogenetic analysis by Maximum Likelihood method (timetree) for NIRVS with similarities to the L protein of Rhabdovirus. A timetree inferred using the RelTime method (Tamura et al. 2012) and the JTT matrix-based model (Jones et al. 1992) is shown. The estimated log likelihood value is -116005.08. A discrete Gamma distribution was used to model evolutionary rate differences among sites (2 categories (+G, parameter = 0.8331)). The rate variation model allowed for some sites to be evolutionarily invariable ([+I], 0.21% sites). The analysis involved 49 amino acid sequences. There were a total of 2319 positions in the final dataset. *(B)* Molecular phylogenetic analysis by Maximum Likelihood method (timetree) for NIRVS with similarities to the G protein of representative Rhabdoviruses (Palatini et al. 2017). Timetree was inferred using the RelTime method (Tamura et al. 2012) and the Whelan and Goldman model (Whelan and Goldman 2001). The estimated log likelihood value is -3719.06. A discrete Gamma distribution was used to model evolutionary rate differences among sites (2 categories (+G, parameter = 2.1095)). The analysis involved 40 amino acid sequences. All positions with less than 95% site coverage were eliminated. That is, fewer than 5% alignment gaps, missing data, and ambiguous bases were allowed at any position. There was a total of 56 positions in the final dataset. *(Cj* Molecular phylogenetic analysis by Maximum Likelihood method (timetree) for NIRVS with similarities to the NS3 protein of Flaviviruses. Timetree was inferred using the RelTime method (Tamura et al. 2012) and the Le_Gascuel_2008 model (Le and Gascuel 2008). The estimated log likelihood value is -6360.35. A discrete Gamma distribution was used to model evolutionary rate differences among sites (2 categories (+G, parameter = 0.8640)). The analysis involved 30 amino acid sequences. All positions containing gaps and missing data were eliminated. There was a total of 180 positions in the final dataset. *(Dj* Molecular phylogenetic analysis by Maximum Likelihood method (timetree) for NIRVS with similarities to the NS5 protein of Flaviviruses. Divergence times for all branching points in the topology were calculated using the Maximum Likelihood method based on the JTT matrix-based model (Jones et al. 1992). The estimated log likelihood value of the topology shown is -26019.44. A discrete Gamma distribution was used to model evolutionary rate differences among sites (2 categories (+G, parameter = 1.0058)). The rate variation model allowed for some sites to be evolutionarily invariable ([+I], 10.15% sites). The tree is drawn to scale, with branch lengths measured in the relative number of substitutions per site. The analysis involved 33 amino acid sequences. There was a total of 984 positions in the final dataset. For all trees, evolutionary analyses were conducted in MEGA7 (Kumar et al. 2016). NIRVS belonging to the core of 20 NIRVS always detected among SSMs are indicated by red boxes. The coding sequences for proteins of the Potato Yellow Dwarft virus (PYDV) were used as outgroup for trees of R-NIRVS, considering PYDV belongs to the highly divergent *Nucleorhabdovirus* genus (Dietzgen et al. 2017). To derive the genealogy of F-NIRVS, outgroups were protein sequences from Tamana Bat Virus (TABV) (de Lamballerie et al. 2002).

**Fig. 5:**
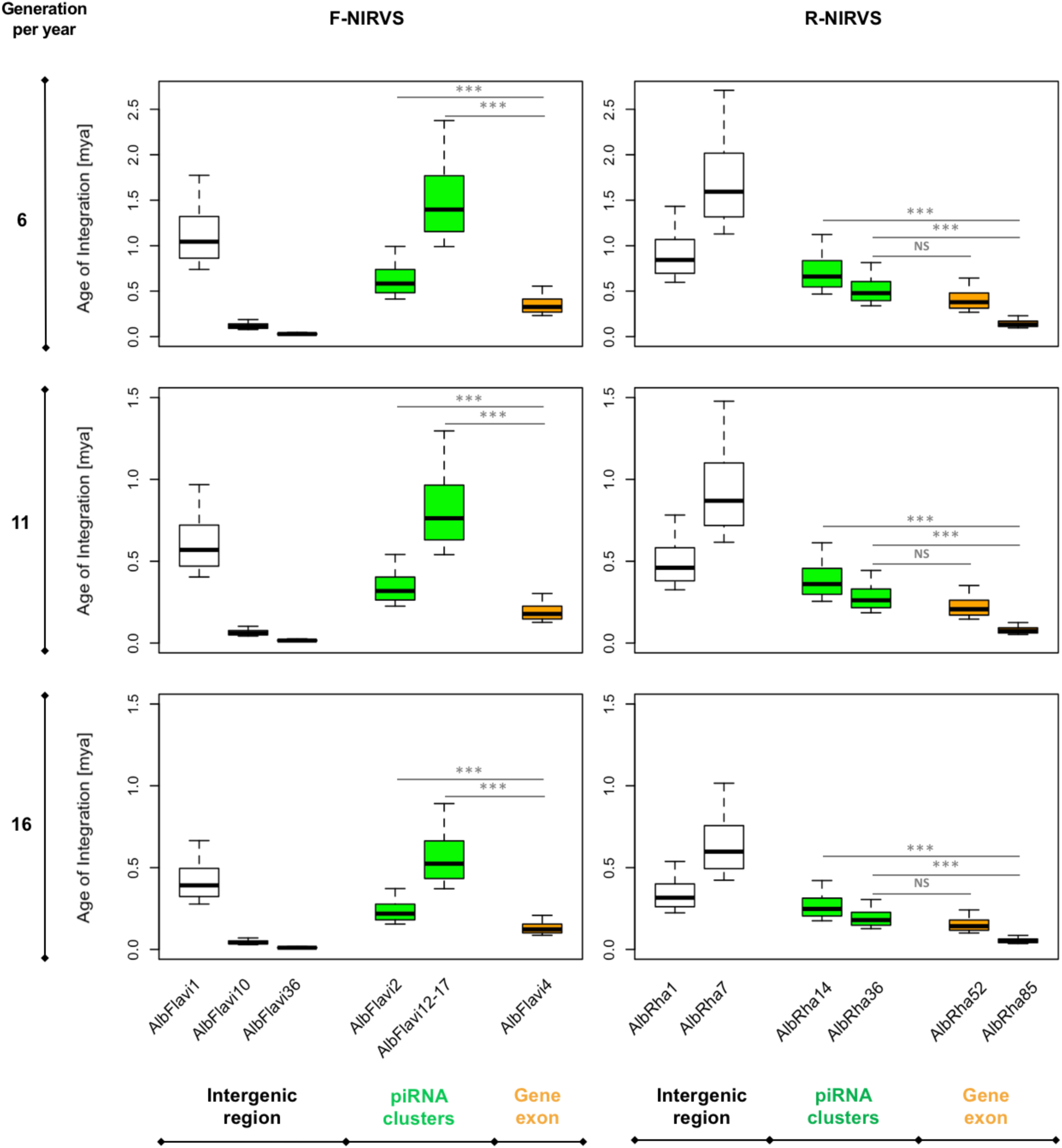
NIRVS integration times. Boxplots showing the integration age for the NIRVS whose variability was studied across five geographic populations. Estimates are based on the *Drosophila* mutation rate, 3.5-8.4×10^-9^ per site per generation (Haag-Liautard et al. 2007; Keightley et al. 2009), and a range of generations per year between 6-16 to include mosquitoes from temperate and tropical regions (Manni et al. 2017). NIRVS of piRNA clusters are statistically older than NIRVS mapping in gene exons (ANOVA, p<0.001).

### NIRVS are heterogeneously polymorphic at the sequence level, with the majority being less variable than slowNevolving genes

The principle that standing genetic variation is not uniformly distributed within a genome, with signals of selection being enriched in specific parts, has been extensively applied to identify regions under positive or negative selection, to test for recent adaptive events and also infer genotype to phenotype associations from polymorphism data (Barrett and Schluter 2007; Charlesworth et al. 2017; Pelissie et al. 2018). On this basis, we selected genes having low and high evolutionary rates in *Ae. albopictus* and we compared their levels of polymorphism (LoP) with that of NIRVS in WGS data from our 16 SSMs. We expanded the analyses to include also genes of the RNAi pathway, for which intraspecific rapid evolution has been observed in *Ae. aegypti* (Bernhardt et al. 2012), and the thirteen genes harboring NIRVS in their coding sequence or UTRs (Palatini et al. 2017).

Estimates of gene evolutionary rates were derived from comparisons of levels of protein sequence divergence within groups of orthologous genes across 27 insect species of the Nematocera sub-order. The first and last 0.1% of groups from the evolutionary rate distribution were selected as slow and fast evolving genes, respectively, and their single-copy orthology status with respect to *Ae. albopictus* was verified. We did not select genes based on their length because of the wide length range of NIRVS, which includes partial viral ORFs of between 151-3206 bp (Palatini et al. 2017). Conserved genes (CGs, slow-evolving) that met the above criteria include genes with hypothetical protein transporter or vesicle-mediated transport activity (*i.e.* AALF003606, AALF014156, AALF014287; AALF014448; AALF004102), structural activity (AALF005886, annotated as tubulin alpha chain), signal transducer activity (AALF026109), protein and DNA binding activity (AALF027761, AALF028431), SUMO transferase activity (AALF020750), the homothorax homeobox encoding gene AALF019476, the tropomyosin invertebrate gene (AALF0082224), the Protein yippee-like (AALF018378) and autophagy (AALF018476). Variable genes (VGs, fast-evolving) include genes with unknown functions (AALF004733; AALF009493; AALF009839, AALF012271, AALF026991, AALF014993, AALF017064, AALF018679), proteolysis functions (AALF010748) a gene associated with transcriptional (AALF022019), DNA-binding (AALF019413, AALF024551), structural (AALF028390) and proteolytic (AALF010877) activities. Median LoP of CGs within mosquitoes of the Foshan strain is 0.0071, a value higher than that observed across 63.3% of the detected NIRVS (supplementary fig. S2). Eleven out of fourteen VGs were more variable than CGs, with seven appearing also statistically more polymorphic than CGs (Kolmogorov-Smirnov test, p< 0.05) (fig. 6A, supplementary table S1A). This result further supports our selection of CGs and VGs. Genes of the RNAi pathway are heterogeneously polymorphic (fig. 6A), with *Ago1* (AALF020776) and *piwi6* (AALF016369) being statistically more polymorphic than CGs; the opposite result was obtained for *piwi 1* and *3* (AALF005499, AALF005498), and *Ago2* (AALF006056) (fig. 6A). LoP of NIRVS is heterogeneous both among SSMs and loci (fig. 6B-C, supplementary table S1B). NIRVS identified within piRNA clusters (Liu et al. 2016) are all less polymorphic than CGs, with the exception of AlbFlavi12-17 that has a median LoP value of 0.0258. This large LoP may be due by the fact that AlbFlavi12-17 is composed of four small viral sequences nested one next to the other (Palatini et al. 2017). Unlike NIRVS from piRNA clusters, NIRVS spanning gene exons are more heterogeneous; three (*i.e.* AlbFlavi34, AlbRha12 and AlbRha52) have LoP values higher than those of CGs, while others (*i.e.* AlbFlavi24, AlbRha28, AlbRha85) are less polymorphic than CGs.

**Fig. 6:**
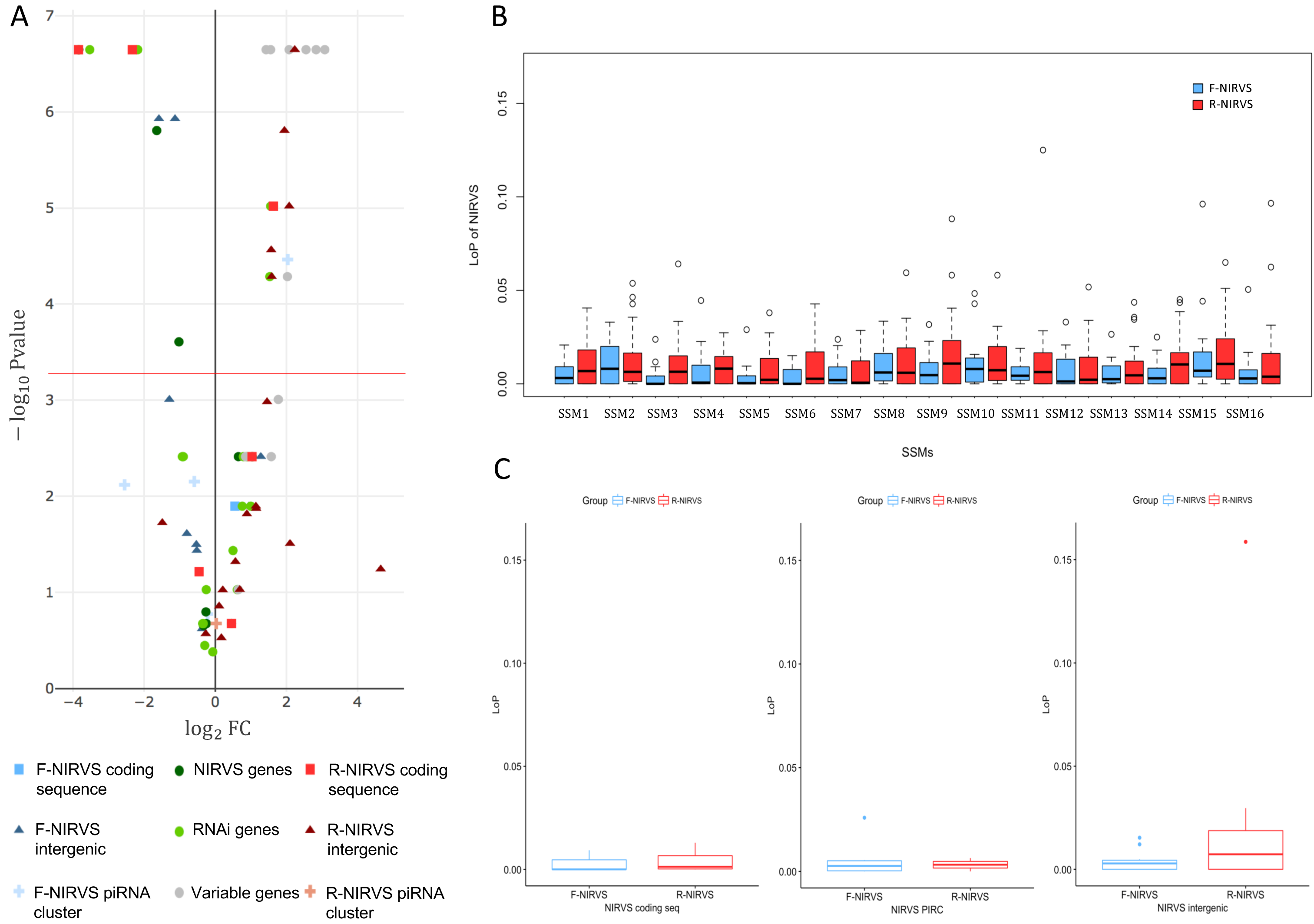
NIRVS polymorphism. *(Aj* Volcano plot showing LoP comparison between CGs and NIRVS, genes encompassing NIRVS (Palatini et al. 2017), genes of the RNAi pathway and VGs. Entities with LoPs statistically different from that of conserved genes are above the red line. Entities on the left side of the panel (log2FC<0) have smaller LoP than conserved genes. The opposite for entities on right side of the panel (log2FC>0). *(Bj* Distribution of LoP values of NIRVS in SSMs are shown for F-NIRVS (blue) and R-NIRVS (red). *(Cj* Distribution of LoP values among loci are shown for F-NIRVS (blue) and R-NIRVS (red).

### SoftNclipped reads support the existence of additional NIRVS variants

We verified the presence of novel NIRVS alleles by investigating soft-clipped reads. Soft-clipped reads support the contiguity of AlbFlavi6 and AlbFlavi7, that were annotated in separated regions of the same contig (fig. 7A). This newly-resolved arrangement revealed the existence of a viral ORF of 1191 bp, corresponding to a partial non-structural protein 5 (NS5) of Flaviviruses. Additionally, soft-clipped reads supported longer than annotated alleles in AlbFlavi10, AlbFlavi2 and AlbRha4 (fig. 7B).

**Fig. 7:**
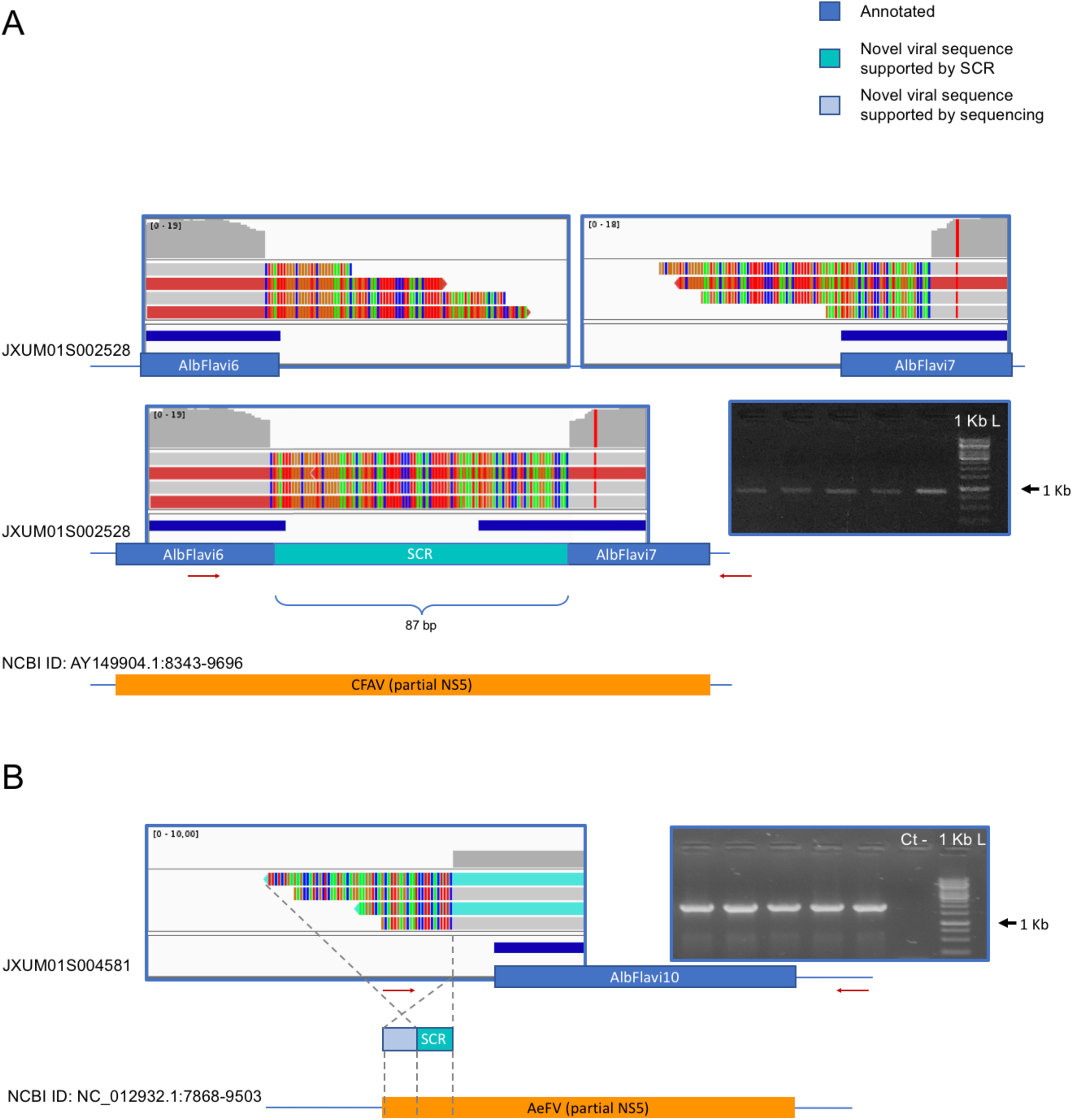
Soft clipped reads (SCR) support novel arrangements and longer than annotated viral integrations. *(A)* AlbFlavi6 and AlbFlavi? are annotated on the same contig, but five thousand bp apart from each other. Soft-clipped reads (in light blue) and PCR experiments support their contiguity, with a unique ORF with similarity to the Flavivirus NS5. *(B)* A sequence of 212 bp extending AlbFlavi10 on its left side was identified investigating SCRs and confirmed by further sequencing. Red arrows indicate primer positions.

### NIRVS identified within coding sequences are expressed

The observed level of polymorphism for AALF020122 with AlbRha52, AALF025780 with AlbRha12, and AALF005432 with AlbFlavi34 is analogous to that of rapidly evolving genes, suggesting co-option for immunity functions (Frank and Feschotte 2017). Because domestication of exogenous sequences is a multi-step process, including persistence, immobilization and stable expression of the newly-acquired sequences besides rapid evolution (Joly-Lopez and Burreau 2018), we analyzed the distribution and expression pattern of these genes. Expression analyses were extended to all other genes harboring NIRVS (AALF025779 with AlbRha9, AALF000476 with AlbRha15, AALF000477 with AlbRha18 and AALF004130 with AlbRha85) that are fixed within the Foshan strain, but have LoP levels comparable to or lower than those of conserved genes (supplementary table S1). AlbFlavi34 had been previously studied and showed to be expressed in pupae and adult males more than in larvae (Palatini et al. 2017). Genes with NIRVS form two groups of paralogs, with similarity to the Rhabdovirus RNA-dependent RNA polymerase (RdRPs) and the nucleocapsid-encoding gene, respectively (table 2). As shown in fig. 8, apart from AALF00477, all other genes are expressed throughout *Ae. albopictus* development with a similar profile, but at different levels. None of the genes showed sex-biased expression or tissue-specific expression in the ovaries, on the contrary highest expression was observed in sugar-and blood-fed females.

**Table 2:** Characteristics of N>Gs.

| **Gene ID** | **NIRVS** | **Viral ORF** | **PfamID** | **Mean LoP** |
| --- | --- | --- | --- | --- |
| AALF000476<sup>a</sup> | AlbRha15 | Rhabdovirus nucleocapsid protein | PF00945 | 0.0086 |
| AALF000477<sup>a</sup> | AlbRha18 | Rhabdovirus nucleocapsid protein | PF00945 | 0.0046 |
| AALF000478<sup>a,c</sup> | AlbRha28 | Rhabdovirus nucleocapsid protein | PF00945 | 0.0004 |
| AALF025780<sup>a</sup> | AlbRha12 | Rhabdovirus nucleocapsid protein | PF00945 | 0.0129 |
| AALF025779<sup>a</sup> | AlbRha9 | Rhabdovirus nucleocapsid protein | PF00945 | 0 |
| AALF004130<sup>b</sup> | AlbRha85 | Rhabdovirus RNA dependent RNA polymerase | PF00946 | 0.0012 |
| AALF020122<sup>b</sup> | AlbRha52 | Rhabdovirus RNA dependent RNA polymerase | PF00946 | 0.0196 |
| AALF005432 | AlbFlavi34 | Flavivirus NS2A, NS2B, NS3 | PF00949, PF00271, PF07652 | 0.0093 |

<sup>a</sup>paralogous genes; <sup>b</sup>paralogous genes; <sup>c</sup>no expression data.

**Fig. 8:**
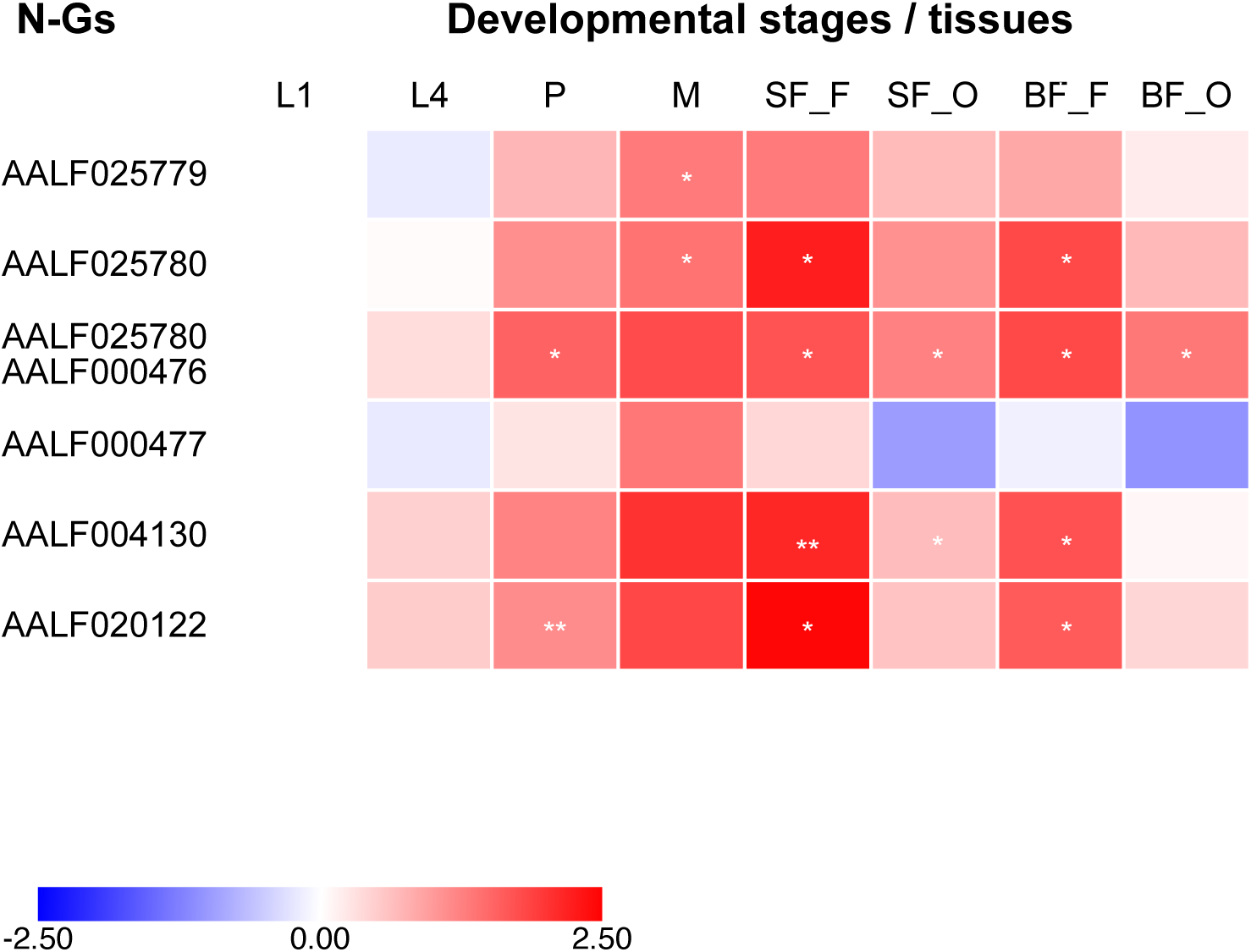
Testing for signs of co-options in six genes harboring NIRVS. *(A)* Heatmap for N-Gs expression profiles across developmental stages and body tissues. Color key expresses the log10- fold change relative to larva 1^st^ instar (calibrator). Asterisks indicate significant differences in transcript abundances (Unpaired 2-tailed t-tests, *P<0.05, **P<0.01).

## Conclusions

Our results represent a large leap forward in the understanding of the composition and structure of the mosquito repeatome. We clearly show that recently-discovered viral integrations are an unappreciated source of genome variability, both structurally and at the sequence level, despite encompassing a limited fraction of the repeatome (27555-36000 bp). This variability is shown not only by the differential distribution of viral integrations in the genome of mosquitoes from different geographic origins and their occurrence in hemizygosity in some mosquitoes, but also by the identification of longer than annotated alleles through soft-clipped reads. A core set of NIRVS, enriched in R-NIRVS, is maintained among all tested mosquitoes. Besides being more widespread, R-NIRVS also appear less variable than F-NIRVS at the sequence level, supporting their older integration into the *Ae. albopictus* genome. Sequence polymorphisms of NIRVS, which was studied in relation to a set of evolutionary-conserved and variable genes, appeared heterogeneous and dependent primarily on their genomic context. NIRVS identified within piRNA clusters were less polymorphic than conserved genes, while among NIRVS encompassing gene exons, three appeared more variable than VGs. Sign of co-option cannot be excluded for two R-NIRVS, AlbRha52 and AlbRha12 with similarity to the RdRPs and nucleocapsid-encoding genes of Rhabdovirus, respectively. RdRPs are ancient enzymes, essential for RNA viruses (de Farias et al. 2017). While the existence of RdRP genes in insects is still debated, cellular RdRP activity has been observed in plants, fungi and *Caenorhabditis elegans* in association with RNA silencing functions (Zong et al. 2009; de Farias et al. 2017; Pinzon et al., unpublished data). An RdRP of viral origin was recently described in a bat species of the *Eptesicus* clade (Horie et al. 2016) and exaptation of a viral nucleocapsid gene was shown in Afrotherians (Kobayashi et al. 2016). On these bases, further experiments to characterize the functions of the *Ae. albopictus* genes AALF020122 and AALF025780 are on-going.

Overall, our analytical approach illustrates the benefits of using genomics and molecular methods to verify evolutionary predictions and identify, within a complex of tens of viral integrations, those that most probably affect mosquito biology. Our data reveal that the impact of viral integrations on mosquito physiology may be more multifaceted than previously thought.

## Material and Methods

### Mosquitoes

Mosquitoes of the Foshan strain have been reared at the insectary of the University of Pavia since 2013 (Palatini et al. 2017). Mosquitoes are reared at 28°C and 70–80% relative humidity with 12/12 h light/dark cycles. Larvae are reared in pans and fed on finely ground fish food (Tetramin, Tetra Werke, Germany). Adults are kept in 30-cm^3^ cages and allowed access to a cotton wick soaked in 0.2 g/ml sucrose as a carbohydrate source. Adult females are blood-fed using a membrane feeding apparatus and commercially available mutton blood.

### Whole Genome Sequencing

The genomes of 16 individual mosquitoes of the Foshan strain were sequenced as previously described (Palatini et al. 2017). Briefly, DNA was extracted from single individuals using the DNeasy Blood and Tissue Kit (Qiagen) and then pooled for Illumina library preparation and sequencing at Polo d’Innovazione di Genomica Genetica e Biologia on a HiSeq2500 platform. Whole genome sequencing data have been deposited at the Sequence Read Archive (SRA) under accession numbers from SAMN09759672 to SAMN09759687.

### Southern Blotting

Genomic DNA (~10 mg) from pools of 10-20 adult mosquitoes of the Foshan strain were digested with restriction enzymes (Thermo Scientific) chosen to specifically target individual F-NIRVS (or groups of F-NIRVS sharing sequence similarity) and separated on a 0.8% agarose gel. DNA was transferred to nylon membranes (Amersham Hybond-N+) and immobilized by UV irradiation. Random-primed DNA probes (Supplementary Material online) were labeled with [a-^32^P] dATP/ml and [a-^32^P] dCTP/ml (3,000 Ci/mmol; 1 Ci- 37 GBq) by using the Megaprime labeling kit (Amersham Pharmacia Biosciences). Hybridizations were carried out at 65°C.

### Real time PCR (qPCR) to test for NIRVS copy number

PCR primers were designed using PRIMER3 (Rozen and Skaletksy 2000) within F-NIRVS to test for their copy number based on real-time PCR (Bubner and Baldwin 2004; Yuan et al. 2007) (Supplementary Material online). Reaction mixtures were prepared containing 10μL QuantiNova Sybr Green PCR Master Mix (Qiagen), 1μL of each 10μM primer, and template DNA diluted in distilled H_2_O up to 20μL total reaction volume. Template genomic DNA used in the reactions was extracted from individual adult mosquitoes following a standard protocol (Baruffi et al. 1995). Real-time PCR reactions were performed in a two-step amplification protocol consisting of 2 minutes at 95°C, followed by 40 cycles of 95°C for 5 sec and 60°C for 10 sec. Reactions were run on a RealPlex Real-Time PCR Detection System (Eppendorf). The single-copy gene *piwi6* (AALF016369) was used as reference. F-NIRVS copy numbers were estimated comparing the relative quantification of NIRVS loci with respect to that of the reference genes using the ⊗Ct method (Pfaffl 2006), after having verified that the efficiencies of PCR reactions with primers for F-NIRVS and the reference gene were the same. Support for using relative quantification without an internal calibrator came from a preliminary test where we cloned one NIRVS (AlbFlavi4) and we verified that estimates of its copy number by absolute vs relative quantification were the same.

### qPCR to estimate NIRVS expression levels

Total RNA was extracted using TRIzol (Life Technologies) from pools including 10-20 mosquitoes at different developmental stages such as larvae, pupae, adult males, sugar-fed females and females sampled 48h after blood feeding. After DNaseI (Sigma Aldrich) treatment, a total of 100 ng of RNA from each pool was used for reverse transcription using the qScript cDNA SuperMix (Quantabio). Expression of the eight genes harboring NIRVS and always detected in Foshan (*i.e.* AALF005432, AALF025780, AALF000476, AALF000477, AALF020122, AALF004130 and AALF025779) was quantified using real-time qPCRs following the protocol described above. Expression values were normalized to mRNA abundance levels of the *Ae. albopictus nap* gene (Reynolds et al. 2012) (Supplementary Material online). The qbase+ software (Biogazelle, Zwijnaarde, Belgium – www.qbaseplus.com) was used to compare expression profiles across samples, and Morpheus (https://software.broadinstitute.org/morpheus) was used to visualize the data.

### Selection of genes with slow and high evolutionary rates

Orthologous genes across 27 insect species within the Nematocera sub-order were identified in OrthoDB v9.1 (Zdobnov et al. 2016). Levels of sequence divergence were computed for each orthologous group as the average of interspecies amino acid identified normalized to the average identity of all interspecies best-reciprocal-hits, computed from pairwise Smith–Waterman alignments of protein sequences (supplementary table S2). We selected the 0.1% of the genes (n=14, number comparable to that of our NIRVS groups) at each tail of the distribution as representative of the conserved and variable categories, the left and right tails respectively. Orthologs of these genes were searched in the *Ae. albopictus* genome (AaloF1 assembly).

### NIRVS in natural populations

PCR primers were designed using PRIMER3 (Rozen and Skaletksy 2000) to test NIRVS polymorphism in *Ae. albopictus* geographic samples (Supplementary Material online). Considering the level of NIRVS sequence similarity, their copy number and heterogeneous presence in Foshan mosquitoes, we selected thirteen NIRVS that gave unambiguous PCR results, have similarities to different viral ORFs and are distributed in different genomic regions including piRNA clusters, intergenic or coding regions. Natural mosquito samples derive from a world-wide collection available at the University of Pavia and previously analyzed with microsatellite markers (Manni et al. 2017). PCR reactions were performed in a final volume of 25 μL using DreamTaq^TM^ Green PCR Master Mix 2x (Thermo Scientific) and the following cycle conditions: 94°C for 3 min, 40 cycles at 94°C for 30 s, 58-60°C for 45 s, 72°C for 1 min, and a final extension at 72°C for 10 min. Amplification products were electrophoresed on 1-1.5% agarose gels and purified using ExoSAP-IT^M^ PCR product Cleanup Reagent (Thermo Scientific). When the NIRVS were detected, at least five amplification products per population per locus were sent to be sequenced by Macrogen (https://dna.macrogen.com/eng/), following the company’s requirements. NIRVS alleles were first scored based on their occurrence in each population and their size. An UPGMA tree was built after 1000 bootstrap resampling of the original data set and the calculation of a matrix of shared allele distances (DAS) by means of the computer program POPULATIONS version 1.2.31 (Langella 1999).

### Estimates of integration time

NIRVS sequences from geographic samples were aligned in Ugene 1.26.1 (Okonechnikov et al. 2012) with MAFFT (Yamada et al. 2016). Default parameters with 5 iterative refinement were applied for the alignment. Alignments were manually curated to verify frameshifts, truncations, deletions, and insertions. All positions including gaps were filter out from the analysis. The following formula was used to estimate the time of integration in years;

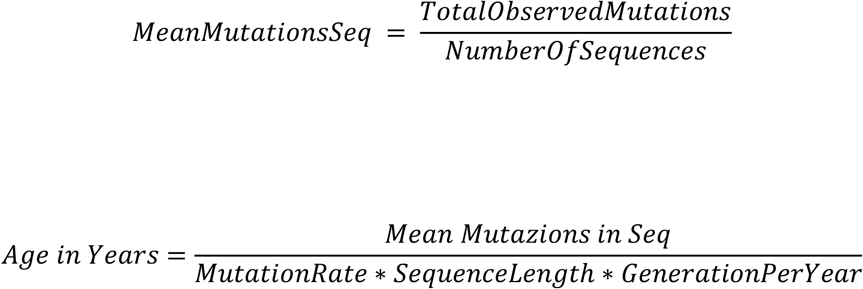

Mutation rate were assumed to be comparable to those of *D. melanogaster* genes in range 3.5-8.4×10^-09^ (Haag-Liautard et al. 2007; Keightley et al. 2009). A range of 4-17 number of generations per year was tested considering mosquitoes of temperate or tropical environments (Manni et al. 2017).

### Phylogenetic inference and timetrees

Deduced NIRVS protein sequences were aligned with subsets of corresponding proteins from Flavivirus and Rhabdovirus genomes using MUSCLE. The timetrees were generated using the RelTime method (Tamura et al. 2011) after having generated the maximum likelihood tree, with 100 bootstrap replicates. Divergence times for all branching points in the topology were calculated using the maximum likelihood method and implementing the best fitted amino acids substitution model. Phylogenies were estimated in MEGA7 (Kumar et al. 2016). Trees were drawn to scale, with branch lengths measured in the relative number of substitutions per site.

### Bioinformatic pipeline to study the polymorphisms of NIRVS

A bioinformatic pipeline was written in Bash and Python languages to test for NIRVS polymorphisms in the WGS data of the 16 SSMs. This pipeline can be used to test the variability of any genomic locus using Illumina paired- end sequencing data and is available upon request. The pipeline is organized into subsequent steps. In the first step, the DepthOfCoverage function of the GATK tool (Mckenna et al. 2010) is used to evaluate the coverage of the region of interest limiting to reads with Phred mapping quality greater than 20. We assumed a NIRVS to be present in a sample when a minimum of 30 consecutive genomic position showed at least 5 reads of depth of coverage. Following read coverage analyses, four different Variant Callers, such as GATK UnifiedGenotyper (Mckenna et al. 2010), Freebayes (Garrison and Marth, unpublished data; https://github.com/ekg/freebayes), Platypus (Rimmer et al. 2016), and Vardict (Lai et al. 2016), are implemented to identify SNPs and INDELs within the regions of interest. The search of SNPs and INDELS by different variant callers allows to increment the pool of variants and reduce the number of false positive. Custom scripts are then used to filter data, retain only variants having allele frequency higher than 0.1 or variants called by two programs.

Follow up statistical analyses were computed and visualized in R studio (R studio Team 2015). The Kolmogorov-Smirnov test was used to test the significance of the difference of LoP distributions of NIRVS, RNAi genes, N-Gs and VGs with respect to that of CGs. CG LoP was the median of the LoPs of the ten tested CGs. The threshold of significance was adjusted with the Bonferroni correction and loci were separated according to the adjusted significance of the test. Results with fold change (FC) different from 0 were visualized in a volcano plot. For each locus, the FC was calculated as the ratio of the median LoP of the locus and that of the CG. The hypergeometric test was applied to test whether the group of NIRVS always identified across SSMs was enriched in 1) F- or R-NIRVS; 2) any viral ORFs; 3) NIRVS shorter or longer than 500bp; 4) NIRVS mapping in exons, piRNA clusters or intergenic regions.

## Acknowledgements

Authors thank Ruth Monica Waghchoure for mosquito maintenance and Lino Ometto for fruitful discussions. This research was funded by a European Research Council Consolidator Grant (ERC-CoG) under the European Union’s Horizon 2020 Programme (Grant Number ERC-CoG 682394) to M.B.; by the Italian Ministry of Education, University and Research FARE-MIUR project R1623HZAH5 to M.B.; and by the Swiss National Science Foundation grant PP00P3_170664 to R.M.W. Authors declare that they do not have conflicts of interest. M.B. and E.P. conceived and designed the study, analyzed data and drafted the manuscript. E.P. and R.M.W. contributed to bioinformatic analyses, analyzed results and revised the manuscript. F.S., F.V., P.L.C. and R.C.L. collected and analyzed molecular data and revised the manuscript. All authors read and approved the final manuscript.

## Supplementary Data

**Supplementary fig. S1.** FGNIRVS Southern blots performed on the genomic DNA of pools of 10G20 Foshan mosquitoes using FGNIRVSGspecific probes (Supplementary Material online). Southern blots corresponding to FGNIRVS predicted to contain partial or complete ORFs of viral proteins (Palatini et al. 2017) are grouped (*i.e.* flaviviral structural E, prM, C and non-structural NS1, NS2a, NS2b, NS3, NS4a, NS4b, NS5).

**Supplementary fig. S2. Distribution of NIRVS, NIRVS genes and RNAi genes based on their LoP levels.** AGF letters indicate six different LoP classes. Grey lines are median LoP values of GCs and VGs. FGNIRVS are blue, RGNIRVS are red, genes encompassing NIRVS are dark green, genes of the RNAi pathway are light green. Within FGNIRVS and RGNIRVS groups, shades of colors are used to highlight NIRVS mapping in exons of annotated genes, piRNA clusters or intergenic regions.

**Supplementary table S1. NIRVS Level of Polymorphisms (LoP) in each SSM and KolmogorovM Smirnov test result.** *(A)* The distribution of LoP of each locus and the distribution of the conserved gene were tested with the KolmogorovGSmirnov test. The threshold of significance was adjusted with the Bonferroni correction and loci were separated according to the adjusted significance of the test (Glog10 0.00048 = 3.32). *(B)* Minimum, mean, median and maximum values of LoP for each NIRVS detected by the bioinformatic pipeline are shown.

**Supplementary table S2. Gene evolutionary rate**.

**Supplementary Material.** *(A)* List of probes used for Southern Blot analyses of F-NIRVS. *(B)* List of primers used for qPCR-based copy number. *(C)* List of primers used for population genetics. *(D)* List of primers used for tracing N-Gs expression profiles. *(E)* Primers used to confirm data from soft-clipped reads.

